# The Euler Characteristic and Topological Phase Transitions in Complex Systems

**DOI:** 10.1101/871632

**Authors:** Edgar C. de Amorim Filho, Rodrigo A. Moreira, Fernando A N Santos

## Abstract

In this work, we use methods and concepts of applied algebraic topology to comprehensively explore the recent idea of topological phase transitions (TPT) in complex systems. TPTs are characterized by the emergence of nontrivial homology groups as a function of a threshold parameter. Under certain conditions, one can identify TPT’s via the zeros of the Euler characteristic or by singularities of the Euler entropy. Recent works provide strong evidence that TPTs can be interpreted as a complex network’s intrinsic fingerprint. This work illustrates this possibility by investigating some classic network and empirical protein interaction networks under a topological perspective. We first investigate TPT in protein-protein interaction networks (PPIN) using methods of topological data analysis for two variants of the Duplication-Divergence model, namely, the totally asymmetric model and the heterodimerization model. We compare our theoretical and computational results to experimental data freely available for gene co-expression networks (GCN) of Saccharomyces cerevisiae, also known as baker’s yeast, as well as of the nematode Caenorhabditis elegans. Supporting our theoretical expectations, we can detect topological phase transitions in both networks obtained according to different similarity measures. Later, we perform numerical simulations of TPTs in four classical network models: the Erdős-Renyi model, the Watts-Strogatz model, the Random Geometric model, and the Barabasi-Albert. Finally, we discuss some perspectives and insights on the topic. Given the universality and wide use of those models across disciplines, our work indicates that TPT permeates a wide range of theoretical and empirical networks.

## 1. Introduction

Topology is the branch of mathematics concerned with finding properties of objects that are conserved under continuous deformations. The topological properties of a system are usually global properties that are independent of its coordinate system or a frame of reference [1]. There is a vast literature in theoretical physics relating phase transitions with rigorously mathematical topological changes in continuous and discrete systems. Research in continuous systems was made using tools of Morse theory [2, 3, 4, 5, 6, 7], while other applications relate to percolation theory [8, 9, 10, 11] on discrete systems. Over the past years, the increasing availability of technology and new experimental methods has led to a massive data production that requires high processing power and the development of new methods and concepts to analyze them. In this context, topological data analysis (TDA) emerges as a promising methodological tool in complex systems [12, 13].

Phase transitions in the context of stochastic topology started with the work of Erdős and Réyni [14], who investigate the problem of tracking the emergence of a giant component in a random graph for a critical probability threshold. Using methods from algebraic topology, one can generalize the basic concept of graphs onto simplicial complexes, i.e., a set of points, edges, triangles, tetrahedron and their n-dimensional analogues, which was later used by Kahle [15] and Linial [16], among others [17] to rigorously extend the giant component transition to a simplicial complex. In this new language, the generalization of the giant component transition constitutes a major change in the distribution of k-dimensional holes, i.e., the emergence of non-trivial homology groups in a simplicial complex, which are characterized by its k-Betti numbers, *β_k_*.

A central metric in this work is the Euler characteristics (EC), which is defined as the alternating sum of the quantities of *k*-dimensional simplices in a simplicial complex. In short, we can compute the Euler characteristic of a complex network by associating it with a simplicial complex. We can identify the (*k* + 1)-cliques, i.e. the all-to-all connected subgraphs with *k* + 1 nodes in a network with its *k*-dimensional simplices, thus obtaining the so-called clique complex. Therefore, if we denote *Cl_k_* as the total number of *k*-cliques in a complex network, the EC *χ* is given by [18, 19, 20]

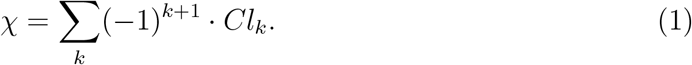

It has been ubiquitously observed that the logarithm density of the Euler characteristic, log(|*χ*|), presents singular behaviour around the thermodynamic phase transition in theoretical models[2, 21], as well as around the zeros of the EC[22, 23, 24, 25, 26]. This intuitively led to the introduction if topological phase transitions in simplicial complexes as the loci of the zeros of the EC [27], where it was observed that the zeros of the Euler characteristic also emerges in the vicinity of the generalization of the Giant component transition [16]. Later, it was observed that the zeros of EC are also associated with the emergence of giant cycles in a simplicial complex [28, 29, 17]. Those transitions can be seen as one possible generalization of percolation transitions to higher-dimensional objects and were observed for the random clique complex of the Erdős-Réyni graph and primarily observed in complex systems for functional brain networks [27]. Later, topological phase transitions were also investigated across different domains, such as financial networks [30], collaboration networks [31], and astrophysics [32], to name a few. Recent developments on high-order interactions on network science [33, 34, 35], persuade us to further explore topological phase transitions in complex systems.

In this work, we explore the topological transition in a variety of models, as well as discuss perspectives and applications in complex systems. As an exemplary case for the ubiquity of phase transitions in complex systems, this work studies the Euler characteristic and its topological phase transitions in two variants of Protein-Protein Interaction Networks (PPIN) models, as well as empirical data and numerical simulations in some classical complex networks.

PPIN are networks in which the nodes are proteins and the edges are the interactions between them. Some topological properties of PPIN have been the subject of many relevant studies [36, 37, 38]. Here, we study the topological phase transitions in two analytical models of PPIN growth: the totally asymmetric and the heterodimerization duplication-divergence models [39, 40].

Moreover, to check whether the topological transitions identified are consistent with real PPIN data, we apply the same approach to data of yeast Gene Co-expression Networks (GCN) freely available online from the project Yeastnet v3 [41] and illustrated in Fig. 1. A GCN is a PPIN in which the edges are based on experimental measures of correlation between the activity of genes over different conditions [42]. We then study the topological phase transitions in the Heterodimerization model [40], a variant of the Duplication model, and compared our results with the phase transitions displayed in the *C. elegans* database [43].

**Figure 1:**
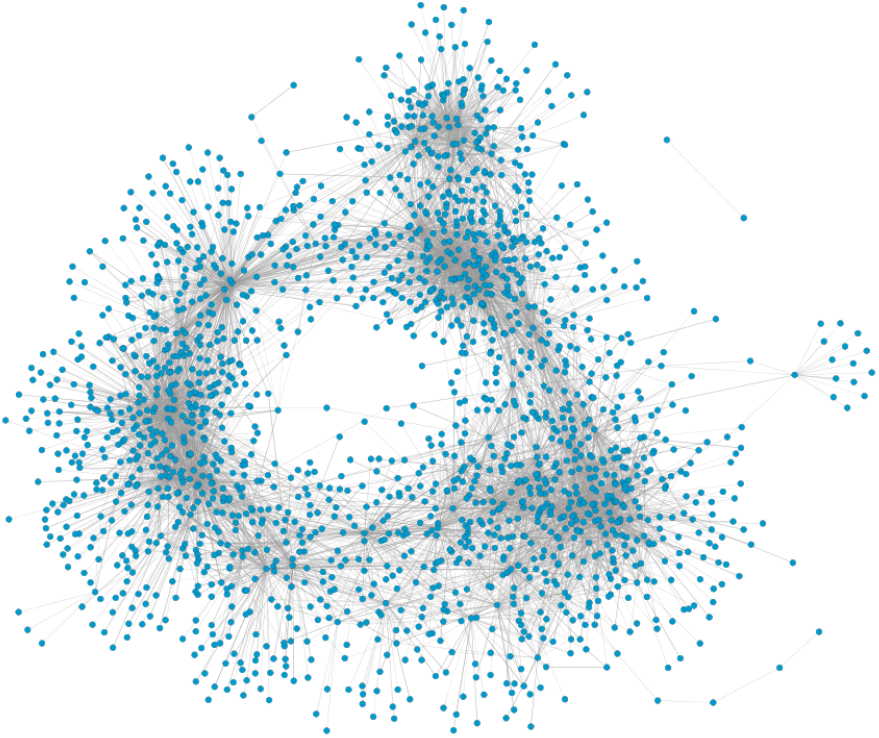
Network representation of gene co-expression network for the yeast using Cytoscape [44]. The presented yeast network has 1, 893 nodes, 7, 955 links and it was obtained from Yeastnet v3 [41].

Then, we move towards the emergence of topological phase transitions in four classical network models that are widely used for network analysis across research fields and scientific disciplines: the Erdős-Réyni, the Watts-Strogatz model, the Random Geometric Graph, and the Barabasi-Albert model. Hereby, we seek to illustrate that topological phase transitions permeate complex systems and might become a useful methodology to characterize the structure and intrinsic properties of complex networks.

This manuscript is written as follows: in section 2 we introduce the protein-protein interaction networks (PPIN) and discuss the duplication-divergence models; in section 3 we study phase transitions in PPIN through the totally asymmetric model, in section 4 we explore topological phase transitions in yeast gene co-expression networks (GCN); and in section 5, we mobilize the heterodimerization model and apply it to data on the nematode *C.elegans*. Moving from PPIN to broader theoretical and simulated models, in section 6, we study the topological phase transitions in four classical complex network models. Finally, section 7 offers our concluding remarks.

## 2. From Protein Interaction Networks to Duplication-Divergence Model

Proteins are macromolecules that participate in a vast range of cell functions, such as DNA replication, responses to stimuli or even the transport of molecules. The interaction between proteins represents a central role in almost every cellular process [45]. For instance, understanding how proteins interact within a network can be crucial to advance the identification of cells’ physiology between normal or disease states [46].

Interactions between proteins in a cell can be mathematically represented as a network (or graph), a mathematical structure composed by nodes (or vertices) and edges (or links). In this framework, proteins are the nodes and each pair of interacting nodes are connected by an edge. This representation is often referred to as a Protein-Protein Interaction Network [36].

In [36] it was first observed that PPIN behave similarly to scale-free networks, i.e., that there are a few nodes participating in the majority of cellular functions. Those nodes are connected to many other nodes and have a high degree, the so-called Hubs. In fact, this scale-free property can be observed in a range of other complex systems, like the world wide web or social networks [47]. The emergence of such systems is characterized by continuous growth and a preferential attachment principle: new nodes are always been added to the system and have a higher propensity to attach to nodes with a higher degree, resulting in a power-law degree distribution. In such distribution, the fraction *p*(*k*) of nodes with degree *k* in the network is inversely proportional to some power of *k*. In PPIN, this scale-free topology is reported to be a consequence of gene duplication [48].

Gene duplication is the process that generates new genetic material during molecular evolution. We can assume that duplicated genes produce indistinguishable proteins which, in the network formalism, implies that new proteins will have the same link structure as the original protein. Every time a gene duplicates, the proteins that are linked with the product of this gene have one extra link on the network. Thus, proteins with more links are more likely to have a neighbour to be duplicated.

The discovery of a scale-free property in PPIN gave rise to many models for generating PPIN based on the gene duplication principle [49, 50, 39, 40, 51]. Most of these models rely on divergence, a process in which genes generated by the same ancestral through duplication accumulate independent changes on their genetic profile over time. The so-called Duplication-Divergence (DD) models were first introduced in [49]. The models are based on the observation that PPIN grow after several gene duplication that can occasionally diverge in their functions over time. There are some variants of this model, which include for instance heterodimerization, arbitrary divergence [40] and random mutations [50].

## 3. Totally Asymmetric Duplication-Divergence model

The totally asymmetric model is defined as a duplication-divergence model in which the divergence process is assumed to happen only on the replica of the duplicated node. This model, proposed in [39], is based on the hypothesis that, after duplication, there is a slight chance that the replicate node starts to develop different functions from the original one. In the model, this change is indicated by the deletion of some edges from the replica and is supposed to happen a single time during the growth of the network. The growth of the network according to this model is illustrate in Figure 2 and it is defined as follows: Given a number *N* of proteins and a probability *p* (0 ≤ *p* ≤ 1), the model generates a graph with N nodes starting from a single one, following the presented algorithm:

- Duplication: One node, as well as its edges, is randomly selected to duplicate.
- Divergence: Each duplicated edge that emerges from the replica is activated with a retention probability *p*. The non-activated edges are removed from the network.

**Figure 2:**
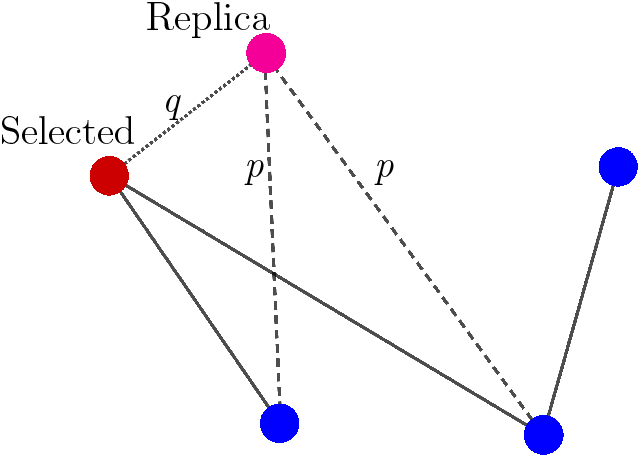
Illustration of a duplication step in PPIN model. For each duplication step a node is selected to be duplicated (red) with its edges. Each new duplicated edge (dashed lines) that goes from the replica (pink) can be activated with independent probability *p* and the non-activated edges are removed. Additionally, for the heterodimerization model, an edge going from the replica to the original node (dotted line) is created with probability *q*.

The duplication step aims to capture genetic replication in a cell, while the divergence step simulates the possibility of a mutation after duplication, which can generate new proteins performing different functions than the original. Some authors [39] consider that any duplication that leads to an isolated node should not be considered. However, since isolated nodes are also considered when computing the Euler characteristic, we kept them to guarantee consistency between theory and experiment. In fact, this approach can be seen as a different perspective in the understanding of gene co-expression data modelling. As we will verify further, experimental data shows isolated nodes for lower levels of co-expression (≪ 1). As we illustrate below, keeping the isolated duplicated nodes will make the modelling more appropriate to match topological phase transitions in the model with the experiments.

For the computation of the Euler characteristic (or equivalently the Euler entropy), observe that the Duplication-Divergence algorithm will only produce bipartite networks, which implies that the networks produced by this model will not have cliques with size 3 or greater. The EC, therefore, will be reduced to the formula *χ* = *N* — *E*, where *N* represents the total number of nodes, and *E* the total number of edges of the network. This simplicity enables us to analytically obtain the expected mean value of the EC in terms of the probability *p* for a graph with *N* nodes, by

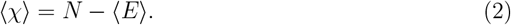

The expected value for the number of edges as function of *p* for a totally asymmetric model with *N* nodes given by:

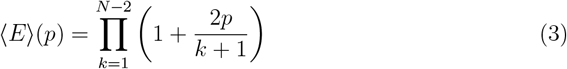

Since the Euler entropy is given by *S_χ_* = log |〈*χ*〉|, we have a singularity on *S_χ_* at the values of p where 〈*χ*〉 = 0, i.e., where

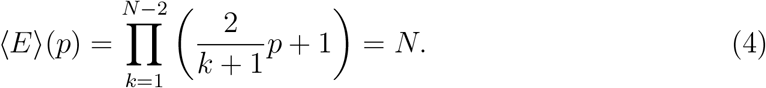

A network that has more edges than nodes certainly has cycles [52]. So, the first topological transition point marks the exact retention probability where it is highly probable that the network has cycles, i.e., in the vicinity of the emergence of a Giant component in the Network [14].

In figure 3, we observe the behaviour of *S_χ_* as a function of the probability *p* of the totally asymmetric model, for a network with 1, 000 nodes. One can observe that there is a single singularity when *p* reaches the critical value of ≈ 0.56 (dashed black line), in the vicinity of the critical probability of the Giant component transition.

**Figure 3:**
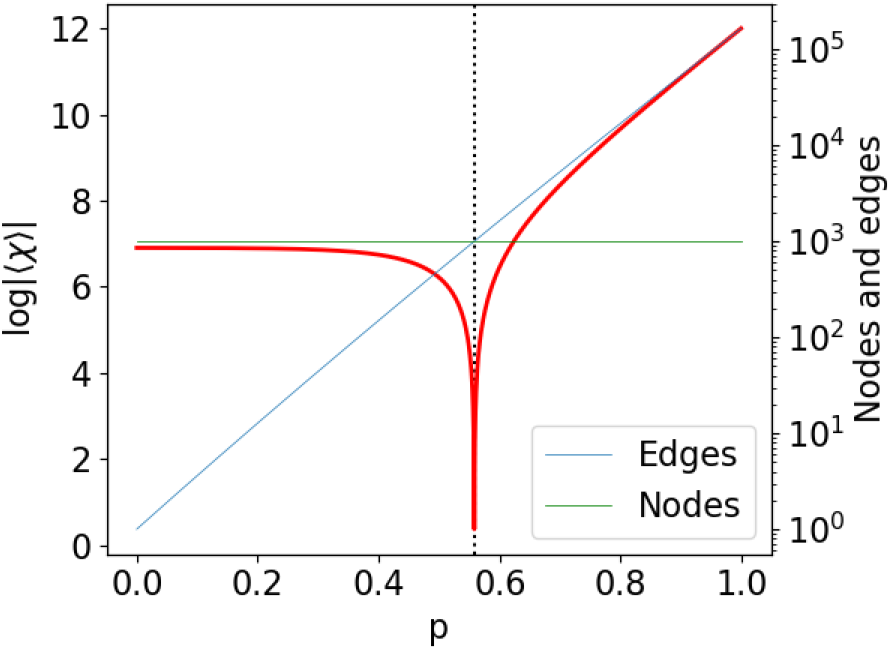
Euler entropy *S_χ_* = log |〈*χ*〉| as function of the retention probability *p* for the Duplication Divergence model with 1, 000 nodes. *S_χ_* was determined by (2). The singularities *S_χ_* → −∞, or the zeros of *χ* locate the topological phase transitions on the duplication divergence model. We put the number of nodes and edges in a logarithmic scale to make visible the change of domain between the number of nodes and the number of edges that occurs at the phase transition’s critical probability.

To characterize this phase transition and understand its implications, we analyze the model through another topological parameter, known as Betti numbers. In general, Betti numbers, indicated by *β_n_*, are defined as follows: *β*_0_ is the total number of connected components. *β*_1_ is the number of cycles on the graph. For *n* > 1 the structure becomes more complex, but roughly speaking, *β_n_* indicates the number of *n*-dimensional “holes” in the network [18, 53].

Figure 4 presents the behaviour of the Betti numbers *β*_0_ and *β*_1_ as functions of *p* compared with the Euler characteristic. Here, we were not able to achieve an analytic expression for the Betti numbers, they were obtained numerically by generating and averaging 1, 000 simulations for each value of *p* (from 0.0 to 1.0 at steps of 10^−2^), 1, 000 nodes each.

**Figure 4:**
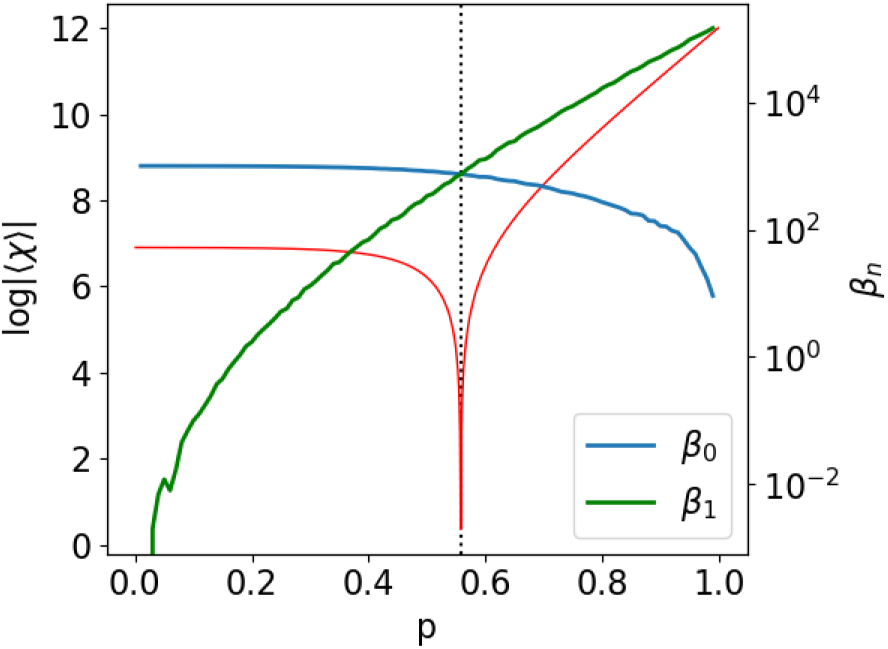
Betti numbers *β*_0_ and *β*_1_ for the Duplication Divergence model as function of the retention probability *p* (*N* = 1,000). At the first phase the network is fragmented, with many components. As *p* increases, *β*_1_ (i.e, the number of cycles) surpasses *β*_0_ when *p* crosses the transition point, indicating a major change in the topology of the DD graph.

Notice that the topological transition occurs in the vicinity of the value of *p_c_* where the Betti curves *β*_0_ and *β*_1_ overlap and, consequently, *χ* = 0, thus confirming the results introduced theoretically in the context of stochastic topology [15, 16, 17]. For *p* < *p_c_* the network is divided into many components because there is a high chance that duplicated nodes do not connect with any neighbour of the original nodes. As *p* increases, the number of components decreases and we have an abundance of cycles in the network. Near *p_c_*, the expected number of cycles is as big as the expected number of components, so there is a high chance that the graph has a cluster with 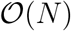 nodes.

Another interpretation for the phase transition obtained from the totally asymmetric model comes from percolation theory. It is a known fact that the first change of sign of the EC in complex networks is related to the appearance of a giant component, i.e., of a connected component in the graph that involves a significant part of the nodes in the system. In the totally asymmetric model, we observe a similar relation, as the topological phase transition obtained is the percolation transition of the system, i.e., the value of the probabilistic parameter that maintains most of the proteins interacting as a whole.

## 4. Gene Co-expression Networks

In a biological context, interactions among proteins can be physical interactions, indicated by the physical contact between them as a result of biochemical events guided by electrostatic forces [54], but also functional interactions. In fact, a group of proteins can perform a common biological function without actually being in direct contact, regulating a biological process or making common use of another molecule [45].

To identify functional interactions, a common method consists in analyzing gene co-expression patterns. This method is based on the assumption that genes with correlated activity produce interacting proteins [55]. Therefore, by mapping the correlation between the activity of genes, we can build the so-called co-expression network (GCN) [45].

A GCN is a PPIN in which the edges are based on experimental measures of correlation between the activity of genes over different conditions. Each node of the network corresponds to a gene. An edge connects a pair of genes if they present similar expression patterns over all the experimental conditions [42]. GCN are often used to identify groups of genes that, over various experimental conditions, displays correlated expression profiles. Based on the “guilt-by-association” principle [56], it is possible to hypothesize that those genes share common functionality. Therefore, the understanding and comparison of topological features across co-expression networks may provide useful information about the strengths (and weaknesses) of the theoretical models used to infer different co-expression networks [57].

We now contrast the theory and numerical simulations presented in the previous section with gene co-expression networks obtained experimentally from data for yeast networks available in [41].

Our aim is to test whether we observe similar topological transitions on real data of GCN, and if such transitions provide us with any useful insights about the structural properties of the network.

Here, we analyzed 48 empirical GCN from *Saccharomyces cerevisiae*, also known as baker’s yeast, available online from the project YeastNet v3 [41]. The whole data set covers around 97% of the yeast coding genome (5, 730 genes). The Networks in this data set have between 800 and 3, 000 nodes and between 7, 000 and 64, 000 links. They were collected through diverse experimental approaches. Each of the data sets analyzed consists of one adjacency matrix for the yeast network. Each node on this network corresponds to a gene. Two genes are connected by an edge if there is a significant co-expression relationship between them. We associate to each edge a weight that corresponds to the absolute value of the correlation between the expression levels of the nodes in the network.

After creating GCN from the co-expression matrices, we proceed with a numerical process called filtration. For a given *ε* ∈ [0,1], this process requires the removal of all the edges with correlation at most 1 — *ε*. For *ε* = 0, for example, we would have an empty network, with all the nodes disconnected. As ε increases, we include the links with a correlation greater than 1 — *ε*. We perform this procedure until *ε* reaches 1 and the network becomes fully connected. In this paper, we used the filtration process to track how the topology of the GCN changes as a function of the correlation between the genes.

In contrast to the results of the previous case, where the EC was written analytically as a function of the probabilistic parameter *p* for the totally asymmetric model, the EC can be obtained numerically as a function of a parameter *ε*, such that *Cl_k_* = *Cl_k_*(*ε*) and consequently *χ* = *χ*(*ε*) in Eq. (1).

As the computation of topological invariants is an NP-complete problem [58], here, we only illustrate data-sets where the computation was feasible, namely, to 20 Yeast networks. Figure 5 shows the Euler entropy as a function of *ε* for the 20 selected networks on Yeastnet’s database. The time scale for the numerical computation of the numbers of *k*-cliques increases exponentially with the size of the network. This occurs because, as *ε* increases, the network becomes denser, and the computation of the cliques become extensive. For this reason, in some of the datasets, we could not compute the Euler characteristic to a threshold where one could detect a topological phase transition. Nonetheless, the Euler entropy averaged over the data sets (blue line) clearly shows the presence of a singularity.

**Figure 5:**
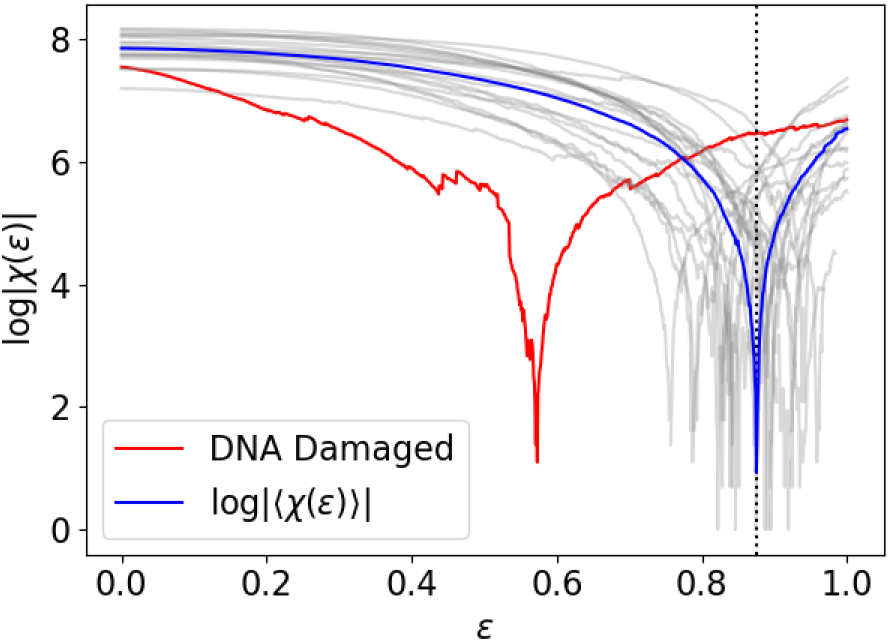
Euler entropy *S_χ_* as a function of the correlation threshold *ε* for 20 co-expression networks from Yeastnet database. Each gray line corresponds to a unique co-expression network, while the blue line is the average of the gray ones. Most of the topological phase transitions were detected on those networks for *ε* in the average critical value *ε* ≈ 0.876, except for the yeast network with DNA damage, where the transition happens for *ε* ≈ 0.573 (red curve). This particular experiment was designed to measure the response to DNA damage in the network [59], and the shift in the critical threshold of the topological transition is apparently sensitive to that response.

Observe that most of the singularities happen near *ε* ≈ 0.876 except for one of the networks, in red, where the singularity happened at *ε* ≈ 0.573. This network comes from an experiment whose goal was to evaluate response to DNA damage due to methylmethane sulfonate and ionizing radiation [59]. In fact, It is known that eukaryotic cells respond to DNA damage by rearranging its cycle and modulating gene expression to ensure efficient DNA repair [59]. Therefore, our analysis suggests that the Euler entropy could be sensible to this reorganization process in the damaged yeast network. The purpose of this empirical example is for illustration of the methodology, and therefore further investigation on the relationship between GCN with DNA damage and percolation transitions will be an important step to establish whether the zeros of the Euler characteristic can be used as a topological bio-marker of such systems, with the potential to detect DNA damage.

In order to provide a deeper characterization of the topological phase transition of GCN, we also calculated the Betti curves *β*_0_ and *β*_1_ for the same yeast networks From a theoretical perspective, a topological phase transition should occur at the vicinity of the zeros of EC and when *β*_0_ ≈ *β*_1_. In Figure 6, we illustrate the averaged Betti curves for the above-mentioned dataset.

**Figure 6:**
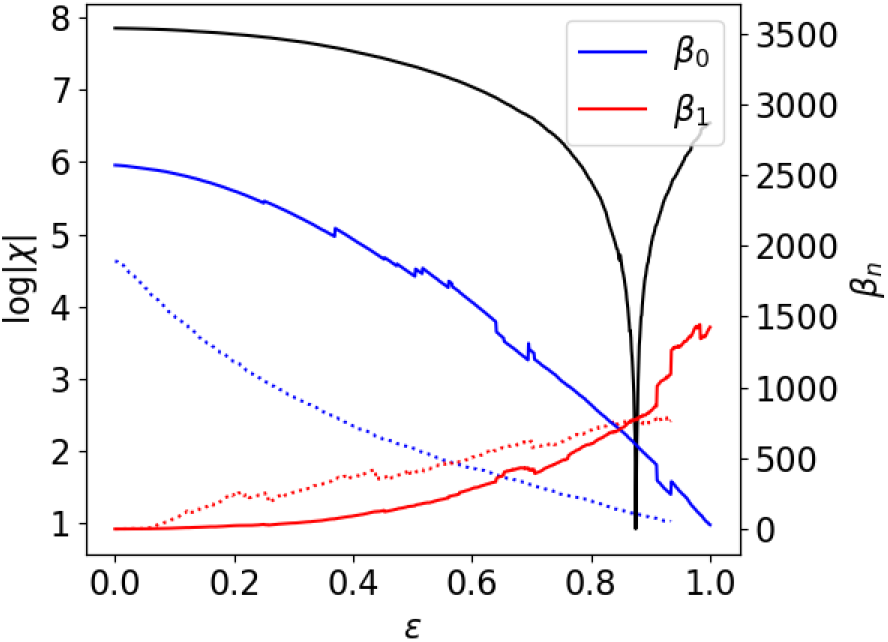
Average Betti curves (*β*_0_ and *β*_1_ as a function of the correlation threshold *ε*). Observe that, as with the totally asymmetric model analyzed previously, the domain of *β*_0_ and *β*_1_ shifts at the vicinity of the singularity of the Euler entropy (black line), i.e., at the zeros of the Euler characteristic. Dotted lines correspond to the Betti numbers of the Yeast network with DNA damage.

We can observe agreement between the threshold transition of *S_χ_* and the threshold where *β*_1_ ≈ *β*_0_, in analogy with the topological transitions reported for theoretical models of random simplicial complexes [16, 60] and for functional brain networks [27].

For values of the correlation threshold below the transition, the network is fragmented in many components, since *β*_0_ is greater than *β*_1_. As *ε* gets closer to the transition, more edges are added to the network, lowering the number of connected components, and changing the network structure to a denser one and with more cycles, i.e., higher *β*_1_. Once the number of connected components approaches 1, i.e., *β*_0_ → 1, almost every new edge has a high probability to create more loops, and the topological transition happens precisely when the number of loops (*β*_1_) is greater than the number of connected components (*β*_0_).

These observations about the behaviour of Betti numbers are compatible with the results observed in [61], and provide some possible interpretation for this shift at the critical point of the topological phase transition. In [61] a linear correlation of *R* = —0.55 was observed between the chance of survival of cancer patients and the number of cycles in the network, also known as the complexity of the cancer PPIN. In fact, the complexity was measured using persistence homology, specifically by the magnitude of the Betti number *β*_1_, which counts the number of one-dimensional cycles in the network. The higher *β*_1_, the higher is the complexity, which implies lower chances of survival [61]. This study provides evidence that, for cancerous cells, the complexity of the PPIN is associated with the health state of the cell.

Now, returning to our analysis of yeast gene co-expression networks, we observed that, for one of the networks, the transition indicated by the Euler characteristic happened at a distinct threshold (≈ 0.573). The analysis of the Betti numbers indicates that this transition is characterized by a shift of dominance between *β*_0_ and *β*_1_, as theoretically described for random simplicial complexes [16]. Thus, during the filtration process, the network corresponding to yeast with DNA damage reaches the topological phase transition earlier than the other data (where the transitions happened near *ε* ≈ 0.876), indicating a rapid appearance of cycles in the network, i.e., the increase of network complexity.

It is important to mention that there are more suitable methods to detect DNA damage, for example, measuring the expression of genes in the DNA repair pathway. But, even though further experimental studies are desired to evaluate the statistical significance of the shift in topological phase transitions for yeast PPIN under DNA damage, our results suggest that phase transitions are indeed intrinsic properties of the system and it seems to be sensitive to DNA damage. Together with recent results in brain networks [27, 62] and with arguments discussed in [61], our results reinforce the hypothesis that topological phase transitions have the potential to be used as intrinsic biomarkers for protein interaction networks more generally.

## 5. Heterodimerization Model and *Caenorhabditis Elegans*

We now move towards a model where protein interactions display more complex dynamics and could potentially capture different empirical data for PPIN. In fact, this model is a variation of the totally asymmetric model discussed in the previous sections. Applied to networks, the model is described as follows: Given a number of nodes *N* and two probabilistic parameters *p* and *q*, the model generates a graph from two nodes and a single edge following the steps below:

- Duplication: One node, as well as its edges, is randomly selected to duplicate.
- Divergence: Each duplicated edge that emerged from the replica is activated with retention probability *p*.
- Heterodimerization: The replicated and the original nodes can connect with probability *q*.

The heterodimerization step mimics the probability that the original node is a dimer, i.e., two molecules joined by bonds that can be either strong or weak. This step is important for clustering and is observed in real PPIN [63]. It is also known that this model produces cliques with similar size and quantity to those observed in some real PPIN [40], contrasting with the totally asymmetric model where we could only observe cliques with small sizes.

As previously defined, the phase transitions are the singularities of the Euler entropy. Given the complexity of the distribution of cliques in these models [40], we were only able to generate these networks numerically, and we further compared them with experimental data. In Figure 7a, we observe the average of the Euler entropy *S_χ_* as a function of the retention probability *p* for a representative network with 1, 000 nodes and different values of *q*. We see that the heterodimerization parameter *q* induces the emergence of further topological phase transitions. These additional topological phase transitions emerge because the heterodimerization step makes possible the appearance of cliques of higher order. The higher the value of q, the more probable it becomes for cliques of larger sizes to appear and, therefore, higher is the like-hood of further topological phase transitions. In this context, it is important to note that [64] reports the abundance of large cliques in real PPIN data, which makes the heterodimerization model more compatible with empirical data.

**Figure 7:**
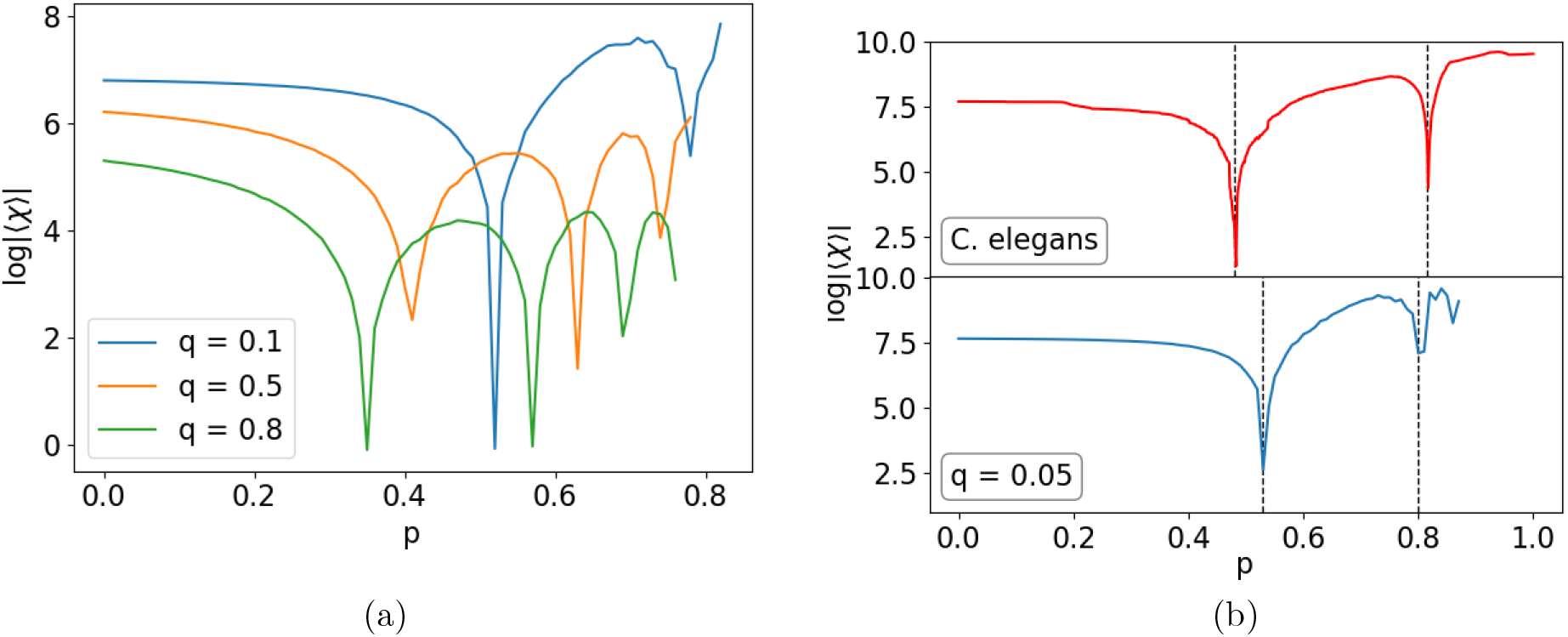
Euler entropy of the Duplication-Divergence model with Heterodimerization. (a) Average of the Euler entropy as function of the retention probability *p* for different values of *q*. Note that, in contrast to the previous model, there are more transitions as we increase the value of *q*. This is because the heterodimerization step makes it possible for cliques of different sizes to appear, as observed in real PPIN [40]. (b) (top) Euler entropy of PPIN from *C. elegans* in contrast with an illustrative simulation for the heterodimerization model with *q* = 0.05 (bottom). Both have two singularities.

Given the bigger size, higher number and complex distribution of the high dimensional cliques in this model, we were not able to achieve an analytic expression for the Euler entropy. Also, since the network becomes denser due to the heterodimerization step, we did not compute the Betti numbers as we did for the totally asymmetric model. Nevertheless, to reinforce the significance of our analysis, we compare our numerical simulations to experimental data for PPIN which, in fact, presents a similar phase transitions profile.

In Figure 7b (top) we illustrate the Euler entropy of a PPIN for the *C. elegans*. The data for this nematode was obtained from the Wormnet v.3 database [43]. For this network, which consists of 2,219 genes and 53,683 links, each link was inferred by analysis of bacterial and archaeal orthologs, i.e., homologous gene sequences of bacteria and archaea related to *C. elegans* by linear descent.

Differently from the data of yeast GCN presented previously, the Euler entropy of this network presents two singularities at the vicinity of *ε*_1_ = 0.48 and *ε*_2_ = 0.82. This behaviour could not be observed if we would consider the growth of a PPIN through duplication and divergence only. However, with the heterodimerization model, we were able to achieve a similar profile of the Euler entropy by simply setting the representative parameters, as can be seen at the bottom of Figure 8b, which shows the expected Euler entropy as a function of the retention probability *p*, averaged over 1, 000 simulations for each value of *p* (ranging from 0 to 1 with steps of 10^-2^). For this simulation, we set the number of nodes at 2, 219 so that we can compare it with the *C. elegans* empirical data. We also set the value of the heterodimerization probability *q* to 0.05. Even though this value was only set for comparison purposes, not intending to perform an optimal fit to the empirical data, it is within the same order of magnitude of the value of *q* = 0.03 used in [40], where the authors managed to achieve a good approximation for the clique distribution across the PPIN considered.

**Figure 8:**
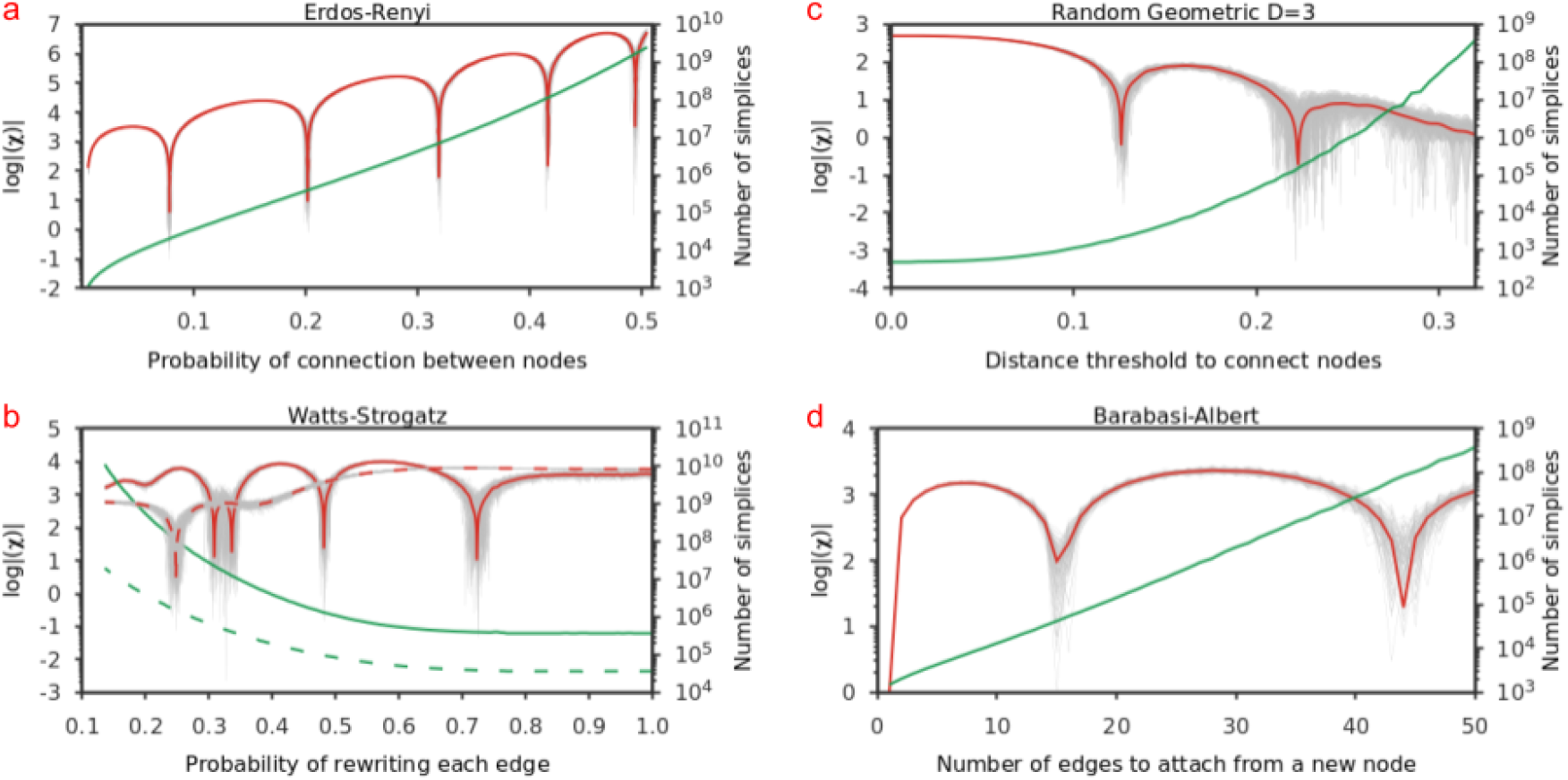
Topological phase transition for classic network models:(a) the Erdős-Rényi model for 0 ≤ *p* ≤ 0.5. (b) the Watts-Strogatz small-world model, for two values of first neighbors connections *k*, namely *k* = 100 (continuous lines), and *k* = 50 (dashed lines). (c) The Random geometric model in *D* = 3 as well as (d) the Barabasi-Albert model. The averages were obtained from 100 realizations for each parameter, green lines are the number of simplices found and red lines the Euler entropy, and gray lines represent individual realizations for each network.

Even though the Duplication-Divergence was enough to understand the topological properties of the yeast networks in the previous sections, it is important to mention that dimerization also happens during the growth of such networks. But this process is not frequent enough to produce significant quantities of big cliques, which would result into new topological phase transitions. This analysis suggests that during the growth of the *C. elegans* network, dimerization happened more frequently, in a way that the Duplication-Divergence model could not reproduce such topological properties. Therefore, the topological phase transitions displayed by the heterodimerization model are more appropriate for this specific empirical network.

Our theoretical and empirical analysis of complex biological networks reinforces the strong evidence that the zeros of the Euler characteristic can be interpreted as an intrinsic fingerprint of a given network. This suggests that topological phase transitions have the potential to be a critical tool to characterize complex systems more generally.

## 6. Topological Phase Transition in Classical Network models

In the remainder of this paper, we illustrate how commonly used complex networks models also displays topological phase transitions. In the previous sections, we showed that topological phase transitions were associated with the log |〈*χ*〉| of the protein-protein interaction network using the Duplication-Divergence and the Heterodimerization models [49], which were subsequently compared against empirical data. Here we investigate the TPT in the Watts-Strogatz model, the Random Geometric model, and the Barabasi-Albert model. Together, these models capture a wide range of phenomena in network science across numerous fields, such as small-world property, percolation transitions, preferential attachment, to name a few.

Once we got the analytical and computational realization of a topological transition in the previous section, now we will illustrate how commonly used complex networks models can be studied using the zeros of Eq. (1), the Euler characteristic *χ*. From a computational perspective, the topological transitions require the use of an ensemble with many replicas for each set of parameters in each model. In this section, we fixed an amount of 100 replicas per system studied. Notice that the number of simplices increases exponentially with the addition of edges to the network. Therefore, without limiting the size of the clique in our computation, we needed to find over 10^11^ simplices per model analysed (see Fig. 8), which is obviously a non-trivial computational problem.

The observation of topological transitions in such models is, therefore, a critical step towards the use of such methodology in complex systems more generally. Moreover, the topological transitions described here are of great interest for a better understanding of high order interactions in complex systems [65, 34, 66, 67, 68, 69, 70].

### TPT in the Erdős-Réyni model

In the Erdős-Réyni network, any two nodes are connected by a linking edge with independent probability *p*. The simplicial complex of the Erdős-Réyni graph, in which all cliques are faces, was investigated rigorously in [15, 27] for fixed *N* nodes as a function of the probability parameter *p* ∈ [0,1]. If *p* = 0, there are no nodes connected (empty graph), whereas for *p* = 1, we have a complete graph. In figure 8 (a), we illustrate the TPT of the ER networks for *N* = 500 nodes and 0 ≤ *p* ≤ 0.5. We can see several singularities in the Euler entropy associated with the many zeros of the mean Euler characteristic. The averages were obtained from 100 realizations for each parameter, green lines are the number of simplices found and red lines the Euler entropy, and grey lines represent individual realizations for each network.

### TPT in the Watts-Strogatz model

To bridge the empirical analysis of protein structures and the discussion of theoretical network models, we now explore the classical Watts-Strogatz network, a small-world model that can also be used to study PPIN [71, 72, 73].

The small-world model is constructed from a ring with *n* nodes, and each node is connected with its *k* first neighbours for *k* even, or *k* — 1 neighbours if *k* is odd. After that, each edge is randomly rewired with probability *p* ∈ [0.0,1.0]. Once these networks are built numerically, we can study the topological transitions displayed by this network as a function of the rewiring probability. In fact, we show in Figure 8 (b) that a small-world network of *n* = 500 also displays the TPT observed in the previous section.

We can proceed with a more detailed inspection of the Watts-Strogatz model by changing the number *k* of first neighbours. For *k* = 50, Fig. 8 shows only one transition. Increasing the number of first neighbours to *k* = 100, we can at least four transitions. Observe that real-world data sometimes have few nodes to be analyzed, and the use of 500 nodes and the ability to differentiate among different topologies (as observed in the previous section) is a clear strength of the present methodology.

### TPT in the Random Geometric model

Once studied through a small-world model, we can go further and, in a second step, investigate the topological transitions of the Random Geometric Graph model in three dimensions [74]. This model is generated by using the number of nodes *n* and a distance *r* as inputs. First, one chooses *n* nodes uniformly at random in the unit cube, which are subsequently connected by an edge if the distance between them is lower than or equal to *r*. We can then study the topological transitions in this network as a function of the distance *r*.

We show the Euler entropy and its topological phase transitions for the Random Geometric model in Figure 8 (c) for *n* = 500 nodes averaged from an ensemble of 100 replicas for each set of parameters. The zeros of EC also correspond to changes in the signs of *χ*, which can be understood as changes in the sign of the mean curvature of the nodes [27] as described by the Gauss-Bonnet theorem for networks [19]. This interpretation can be important when we want to investigate topological changes in network models whose filtration parameter is not necessarily continuous, as we will see below.

### TPT in the Barabasi-Albert model

Lastly, we illustrate the topological changes in the Barabasi-Albert network model [75]. This model is known for introducing random scale-free networks using a preferential attachment mechanism. The network starts with few nodes and no edges and a number of edges attached to the new node with probability proportional to the node’s degree. In contrast to the three previous network models studied in this section, the Barabasi-Albert model has two integers parameters, namely, the maximum number of nodes *n* and the number of edges *m* to attach to a new node to existing nodes.

We can analyse the topological transitions in the Barabasi-Albert network for a fixed *n* = 500 as a function of *m*. However, given the discrete nature of the control parameter *m* of this network model, we were only able to infer the topological phase transitions via changes in the signal of the Euler characteristic, which tracks the changes in the curvature of the nodes via the Gauss-Bonnet theorem for networks [19]. Figure 8 (d) shows three zeros of the Euler characteristics as a function of the number of edges attached using an ensemble of 100 replicas. Similarly to the Random Geometric Graph, the transitions are revealed by a sign change of *χ*. Our inferred results for the first transition, *m* = 2, is in agreement with the percolation transition for the Barabasi-Albert model [76]. However, little is known regarding topological changes in the Barabasi Albert model for higher values of *m*. For this particular model, one interesting analogy could be made regarding the relationship between Lyapunov exponents, curvature and phase transitions known in statistical mechanics [77, 78].

In fact, Lyapunov stability theory was adapted to the context of networks [79], and particularly to the Barabasi-Albert model [80], where the authors studied which values of *m* anomalous events occur in this network. We noticed that some of the critical intervals for *m* in Ref. [80] that are analogous to the changes in the curvature as a function of *m* observed in our work. Therefore, the connection between them deserves further investigation.

Ultimately, the aim of this section was to illustrate that topological transitions, evidenced by the zeros of the Euler characteristic, are present in four classical network models that are widely used in network analysis across a variety of fields. This reinforces the ubiquity of such topological transitions in complex systems more generally.

## 7. Conclusions and Perspectives

In this work, we investigated the topological phase transitions in a wide range of theoretical and empirical networks. Those transitions are characterized by the zeros of the Euler characteristic at a critical probability or threshold and represent a generalization of the giant component percolation transition introduced by Erdős [14, 16, 15] and recently expanded to data-driven complex networks [27]. The zeros of the Euler characteristic also indicate signal changes in the mean curvature [19] and the emergence of a giant *k*-cycle in a simplicial complex [28].

We first focused our attention on protein-protein interaction networks, particularly to networks generated by the totally asymmetric duplication-divergence model. We verified that such transitions correspond to the emergence of a giant component in the network, as observed in yeast and gene co-expression networks datasets [41].

Our results give strong support to the hypothesis that percolation transitions are topological bio-markers of a network [27, 62]. In fact, by analyzing a network for yeast under DNA damage [59], we found that the transition point shifted to a much lower value compared to the interval in which the topological phase transitions of the other yeast network datasets took place. Although this result deserves further investigation, we provide evidence that the zeros of the Euler characteristic can be seen as suitable topological invariants to distinguish macroscopic properties of the yeast networks.

More generally, in order to further explore topological phase transitions in complex systems, we extended our analysis to three classical network models, namely: the Watts-Strogatz model [71], the Random Geometric model [74], and the Barabasi-Albert model [75]. We showed that topological transitions permeate these models which are widely used across disciplines, and some of those transitions were not observed before by the analysis of classical networks using standard methodologies.

Then, we explored topological phase transitions in the *C. elegans* PPIN. In contrast to the yeast networks, the same approach gave rise to two topological phase transitions, indicating that the totally asymmetric model is insufficient to capture the topological aspects of the *C. elegans* PPIN data properly. This suggests that there is another network process at play that leads to the PPIN growth. One possibility could be the presence of dimmers, i.e., two nodes that are parts of the same protein (or gene) that are connected. To test this hypothesis, we computed the expected Euler entropy of the heterodimerization duplication-divergence model and observed that, for an illustrative set of parameters, we could obtain an Euler entropy profile similar to the one observed for the *C. elegans* network. Notice that, in order to match our theory and numerical simulations to the experimental data, we considered that the nodes that got isolated after duplication were kept in the network.

It is important to emphasize that other models suggest different processes for PPIN growth, implying that our analysis did not exhaust the possibilities of models considered for PPIN. In [40] for example, the authors present an arbitrary divergence model in which they could replicate with high confidence the clique distribution observed for PPIN data.

Other models simulate different aspects of the growth of a PPIN [50, 51, 40] and the analysis of such models through the lenses of topological phase transitions and the Euler characteristic would give us precious information about PPIN.

We hope that the overview of TPT reported in this work illustrates the rapid development of phase transitions using algebraic topology methods, especially stochastic topology, in complex systems. There are numerous directions for future works on connecting the results reported here with structure and dynamics in high order networks and complex systems. In short, this work contributes to a better understanding of the ubiquity of topological transitions on real-world data and complex systems, and the possibility of the use of algebraic topological methods to characterize those systems using high-order topological structures.

## 8. Acknowledgements

This work is funded by CAPES, CNPq, and FACEPE (Brazilian Agencies). F. A. N. Santos would like to express his gratitude for the useful discussions with Linda Douw, Mauricio D. Coutinho-Filho, Ernesto P. Raposo, Mauro Coppeli, James Charles Phillips, Stefan Boettcher, Bertrand Berche, Katharina Natter and Fernando Moraes during the development of this work. E. C. de Amorim Filho would like to thank V. A. Caldas and T. Shimizu for the learn-full visiting period at AMOLF. We would like to acknowledge FACEPE Pronex APQ-0602-105/14 and FACEPE-Fulbright ARC-0103-1.01/17.

## 9. Appendix

### Proof of Equation 3

Randomly selecting a node *v* on a network with *N* — 1 vertices, the expected increase in the number of edges after the divergence steps is

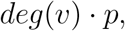

where *deg(v)* is indicating the number of nodes linked to *v* by an edge. So, if we want the expected increase after the duplication and divergence steps, we just need to average over all nodes, giving us

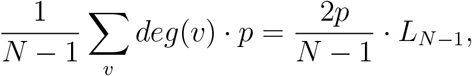

where *L*_*N*-1_ is the number of edges before duplication and divergence steps. Adding that to the edges we had before duplicating, we have that

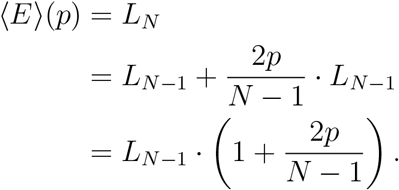

Now, remember that the algorithm starts with a graph with two nodes and a single edge, meaning that *L*_2_ = 1. So, using recursion, we can conclude that

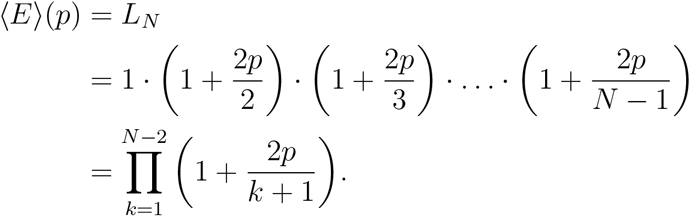

## References

[1] Nakahara M 2003 Geometry, topology and physics (CRC press)

[2] Pettini M 2007 Geometry and topology in Hamiltonian dynamics and statistical mechanics (New York: Springer-Verlag New York) ISBN 978-0-387-30892-0

[3] Kastner M 2008 Reviews of Modern Physics 80 167–187 URL https://journals.aps.org/rmp/pdf/10.1103/RevModPhys.80.167

[4] Santos F A N, da Silva L C B and Coutinho-Filho M D 2017 Journal of Statistical Mechanics: Theory and Experiment 2017 013202 ISSN 1742-5468 URL http://stacks.iop.org/1742-5468/2017/i=1/a=013202?key=crossref.6284bdda4a6c2e768f85850c94d27803

[5] Buchanan M 2008 Nature Physics 4 5–5 ISSN 1745-2473 URL http://www.nature.com/articles/nphys819

[6] Gori M, Franzosi R and Pettini M 2018 Journal of Statistical Mechanics: Theory and Experiment 2018 ISSN 17425468

[7] Kastner M and Schnetz O 2008 Physical Review Letters 100 ISSN 00319007

[8] Speidel L, Harrington H A, Chapman S J and Porter M A 2018 Phys. Rev. E 98(1) 012318 URL https://link.aps.org/doi/10.1103/PhysRevE.98.012318

[9] Okun B L 1990 Journal of Statistical Physics 59 523–527 ISSN 0022-4715 URL http://link.springer.com/10.1007/BF01015581

[10] Mecke K R and Wagner H 1991 Journal of Statistical Physics 64 843–850 ISSN 00224715

[11] Neher R A, Mecke K and Wagner H 2008 Journal of Statistical Mechanics: Theory and Experiment 2008 P01011 ISSN 1742-5468 URL http://stacks.iop.org/1742-5468/2008/i=01/a=P01011?key=crossref.49687f21268502da4a5a8e6780e62649

[12] Edelsbrunner H, Harer J et al. 2008 Contemporary mathematics 453 257–282

[13] Carlsson G 2009 Bulletin of the American Mathematical Society 46 255–308 ISSN 0273-0979 URL http://www.ams.org/journal-getitem?pii=S0273-0979-09-01249-X

[14] Erdos P and Rényi A 1959 Publicationes Mathematicae (Debrecen) 6 290–297 URL http://snap.stanford.edu/class/cs224w-readings/erdos59random.pdf

[15] Kahle M 2013 arXiv e-prints URL http://arxiv.org/abs/1301.7165

[16] Linial N and Peled Y 2016 Annals of Mathematics 184 745–773 ISSN 0003-486X URL http://annals.math.princeton.edu/2016/184-3/p03

[17] Bobrowski O and Krioukov D 2021 arXiv preprint arXiv:2105.12914

[18] Edelsbrunner H and Harer J J 2010 Computational topology: an introduction (American Mathematical Society) ISBN 9780821849255

[19] Knill O 2011 arXiv e-prints URL http://arxiv.org/abs/1111.5395

[20] Wu Z, Menichetti G, Rahmede C and Bianconi G 2015 Scientific reports 5 1–12

[21] Santos F A and Coutinho-Filho M D 2009 Physical Review E - Statistical, Nonlinear, and Soft Matter Physics ISSN 15393755

[22] Gandolfo D 2005 Physica A: Statistical Mechanics and its Applications 358 22–29

[23] Blanchard P, Dobrovolny C, Gandolfo D and Ruiz J 2006 Journal of Statistical Mechanics: Theory and Experiment 2006 P03011–P03011 ISSN 1742-5468

[24] Rehn J A, Santos F A and Coutinho-Filho M D 2012 Brazilian Journal of Physics 42 410–421 ISSN 01039733

[25] Blanchard P, Fortunato S and Gandolfo D 2002 Nuclear Physics B 644 495–508 ISSN 0550-3213 URL https://www.sciencedirect.com/science/article/pii/S0550321302006818

[26] Blanchard P, Gandolfo D, Ruiz J and Shlosman S 2003 Markov Processes and Related Fields 9 523–545 URL http://citeseerx.ist.psu.edu/viewdoc/download?doi=10.1.1.583.4298&rep=rep1&type=pdf

[27] Santos FAN, Raposo E P, Coutinho-Filho M D, Copelli M, Stam C J and Douw L 2019 Physical Review E 100 032414 ISSN 2470-0045 URL https://link.aps.org/doi/10.1103/PhysRevE.100.032414

[28] Bobrowski O and Skraba P 2020 Physical Review E 101 032304

[29] Bobrowski O and Skraba P 2020 arXiv preprint arXiv:2005.14011

[30] Gidea M 2017 Topological Data Analysis of Critical Transitions in Financial Networks (Springer, Cham) pp 47–59 URL http://link.springer.com/10.1007/978-3-319-55471-65

[31] Lee Y, Lee J, Oh S M, Lee D and Kahng B 2021 Chaos: An Interdisciplinary Journal of Nonlinear Science 31 041102

[32] Giri S K and Mellema G 2021 Monthly Notices of the Royal Astronomical Society 505 1863–1877

[33] Battiston F, Cencetti G, Iacopini I, Latora V, Lucas M, Patania A, Young J G and Petri G 2020 Physics Reports

[34] Millan A P, Torres J J and Bianconi G 2019 arXiv preprint arXiv:1912.04405

[35] Bick C, Gross E, Harrington H A and Schaub M T 2021 arXiv preprint arXiv:2104.11329

[36] Jeong H, Mason S P, Barabasi A L and Oltvai Z N 2001 Nature 411 41–42 ISSN 0028-0836 URL http://www.nature.com/articles/35075138

[37] Yook S H, Oltvai Z N and Barabasi A L 2004 PROTEOMICS 4 928–942 ISSN 1615-9853 URL http://doi.wiley.com/10.1002/pmic.200300636

[38] Maslov S and Sneppen K 2002 Science (New York, N.Y.) 296 910–3 ISSN 1095-9203 URL http://www.ncbi.nlm.nih.gov/pubmed/11988575

[39] Ispolatov I, Krapivsky P L and Yuryev A 2005 Physical review. E, Statistical, nonlinear, and soft matter physics 71 061911 ISSN 1539-3755 URL http://www.ncbi.nlm.nih.gov/pubmed/16089769 http://www.pubmedcentral.nih.gov/articlerender.fcgi?artid=PMC2092385

[40] Ispolatov I, Krapivsky P L, Mazo I and Yuryev A 2005 New Journal of Physics 7 145–145 ISSN 1367-2630 URL http://stacks.iop.org/1367-2630/7/i=1/a=145?key=crossref.9173be6c5c84f98aff1f2bcd76806bcc

[41] Kim H, Shin J, Kim E, Kim H, Hwang S, Shim J E and Lee I 2014 Nucleic Acids Research ISSN 03051048

[42] Petereit J, Harris F C and Schlauch K 2015 petal: A novel co-expression network modeling system 2015 IEEE International Conference on Bioinformatics and Biomedicine (BIBM) (IEEE) pp 234–241 ISBN 978-1-4673-6799-8 URL http://ieeexplore.ieee.org/document/7359686/

[43] Cho A, Shin J, Hwang S, Kim C, Shim H, Kim H, Kim H and Lee I 2014 Nucleic Acids Research 42 W76–W82 ISSN 1362-4962 URL http://academic.oup.com/nar/article/42/W1/W76/2435686/WormNet-v3-a-networkassisted-hypothesisgener

[44] Shannon P, Markiel A, Ozier O, Baliga N, Wang J, Ramage D, Amin N, Schwikowski B and Ideker T 2003 Genome Research 13 2498–2504

[45] Vella D, Zoppis I, Mauri G, Mauri P and Silvestre D D 2017 EURASIP Journal on Bioinformatics and Systems Biology 2017 6 URL https://bsb-eurasipjournals.springeropen.com/track/pdf/10.1186/s13637-017-0059-z

[46] Vidal M, Cusick M E and Barabási A L 2011 Cell 144 986–98 ISSN 1097-4172 URL http://www.ncbi.nlm.nih.gov/pubmed/21414488 http://www.pubmedcentral.nih.gov/articlerender.fcgi?artid=PMC3102045

[47] Barabasi A L and Pósfai M 2016 Network Science by Albert-László Barabási 1st ed (Cambridge: Cambridge University Press) URL http://networksciencebook.com/

[48] Barabási A L and Oltvai Z N 2004 Nature Reviews Genetics 5 101–113 ISSN 1471-0056 URL http://www.nature.com/articles/nrg1272

[49] Vázquez A, Flammini A, Maritan A and Vespignani A 2003 Complexus 1 38–44 ISSN 1424-8492 URL https://www.karger.com/Article/FullText/67642

[50] Pastor-Satorras R, Smith E and Solé R V 2003 Journal of Theoretical Biology 222 199–210 ISSN 0022-5193 URL https://www.sciencedirect.com/science/article/pii/S0022519303000286

[51] Farid N and Christensen K 2006 New Journal of Physics 8 212–212 ISSN 1367-2630 URL http://stacks.iop.org/1367-2630/8/i=9/a=212?key=crossref.d976187ae5c371558c9abecd75b27f7c

[52] Bollobás B 1998 Modern graph theory (Springer)

[53] Zomorodian A J 2005 Topology for computing (Cambridge University Press) ISBN 9780511546945

[54] De Las Rivas J and Fontanillo C 2010 PLoS computational biology 6 e1000807 ISSN 1553-7358 URL http://www.ncbi.nlm.nih.gov/pubmed/20589078 http://www.pubmedcentral.nih.gov/articlerender.fcgi?artid=PMC2891586

[55] Grigoriev A 2001 Nucleic Acids Research 29 3513–3519 ISSN 13624962 URL https://academic.oup.com/nar/article-lookup/doi/10.1093/nar/29.17.3513

[56] De Smet R and Marchal K 2010 Nature Reviews Microbiology 8 717–729 ISSN 1740-1526 URL http://www.nature.com/articles/nrmicro2419

[57] Xulvi-Brunet R and Li H 2010 Bioinformatics 26 205–214 URL https://academic.oup.com/bioinformatics/article-abstract/26/2/205/209253

[58] Bomze I M, Budinich M, Pardalos P M and Pelillo M 1999 The Maximum Clique Problem Handbook of Combinatorial Optimization (Springer US)

[59] Gasch A P, Huang M, Metzner S, Botstein D, Elledge S J and Brown P O 2001 Molecular Biology of the Cell 12 2987–3003 ISSN 1059-1524 URL http://www.molbiolcell.org/doi/10.1091/mbc.12.10.2987

[60] Toth C D, O’Rourke J and Goodman J E 2017 Handbook of Discrete and Computational Geometry, Third Edition 3rd ed (Chapman and Hall/CRC) ISBN 9781498711395

[61] Benzekry S, Tuszynski J A, Rietman E A and Lakka Klement G 2015 Biology Direct 10 32 ISSN 1745-6150 URL http://www.biologydirect.com/content/10/1/32

[62] Gracia-Tabuenca Z, Díaz-Patiño J C, Arelio I and Alcauter S 2020 Eneuro 7

[63] Ispolatov I, Yuryev A, Mazo I and Maslov S 2005 Nucleic Acids Research 33 3629–3635 ISSN 0305-1048 URL https://academic.oup.com/nar/article-lookup/doi/10.1093/nar/gki678

[64] Spirin V and Mirny L A 2003 Proceedings of the National Academy of Sciences 100 12123–12128 ISSN 0027-8424 (Preprint https://www.pnas.org/content/100/21/12123.full.pdf) URL https://www.pnas.org/content/100/21/12123

[65] Tran Q H, Hasegawa Y et al. 2019 Physical Review E 100 032308

[66] Bianconi G, Kryven I and Ziff R M 2019 Physical Review E 100 062311

[67] Kartun-Giles A P and Bianconi G 2019 Chaos, Solitons & Fractals: X 1 100004

[68] Skardal P S and Arenas A 2019 arXiv preprint arXiv:1909.08057

[69] Neuhäuser L, Mellor A and Lambiotte R 2020 Physical Review E 101 032310

[70] Benson A R, Abebe R, Schaub M T, Jadbabaie A and Kleinberg J 2018 Proceedings of the National Academy of Sciences 115 E11221–E11230

[71] Watts D J and Strogatz S H 1998 Nature 440–442

[72] del Sol A, Fujihashi H and O’Meara P 2005 Bioinformatics 21 1311–1315 ISSN 1367-4803 URL https://academic.oup.com/bioinformatics/article-lookup/doi/10.1093/bioinformatics/bti167

[73] Taylor N R 2013 Computational and Structural Biotechnology Journal 5 e201302006

[74] Penrose M 2003 Random Geometric Graphs Oxford Studies in Probability, 5 (Oxford University Press, USA) ISBN 0198506260 URL http://www.worldcat.org/isbn/0198506260

[75] Barabasi A L and Albert R 1999 Science 286 509–512 ISSN 00368075

[76] Pietsch W 2006 Physical Review E - Statistical, Nonlinear, and Soft Matter Physics 73 ISSN 15393755

[77] Caiani L, Casetti L, Clementi C and Pettini M 1997 Physical Review Letters 79 4361–4364 ISSN 10797114

[78] Argoul F and Arneodo A 1986 Lyapunov exponents and phase transitions in dynamical systems Lyapunov Exponents (Springer Berlin Heidelberg) pp 338–360

[79] Ruiz D and Finke J 2018 Journal of Statistical Physics 172 1127–1146 ISSN 00224715

[80] Ruiz D and Finke J 2019 International Journal of Applied Mathematics and Computer Science 29 363–373 URL https://content.sciendo.com/configurable/contentpage/journals002famcs002f29002f2002farticle-p363.

